# Loss of STARD7 Impairs Mitochondrial Phospholipid Homeostasis and Contributes to Mitochondrial Myopathy

**DOI:** 10.1101/2025.11.27.691062

**Authors:** Yasuhiro Horibata, Tomoki Sato, Masahide Ohyama, Sae Yuyama, Masahiko Itoh, Shinji Miura, Akimitsu Konishi, Hiroyuki Sugimoto

## Abstract

Mitochondria are composed of phospholipid bilayers rich in phosphatidylcholine (PC). StAR-related lipid transfer domain-containing protein 7 (STARD7) functions as a lipid transfer protein that plays a crucial role in maintaining mitochondrial PC homeostasis. In this study, we investigated the physiological role of STARD7 in skeletal muscle using muscle-specific knockout (mKO) mice. STARD7 expression was markedly higher in the soleus, a mitochondria-dense slow-twitch muscle, compared with fast-twitch fibers. Although muscle fibers from mKO mice exhibited no apparent structural abnormalities, their endurance exercise capacity was markedly reduced. RNA-seq analysis revealed suppressed expression of fast-twitch-related genes accompanied by a reduction in fast-twitch fibers. At the mitochondrial level, respiratory chain complexes remained intact, but oxygen consumption was consistently decreased. Targeted lipidomic analysis showed decreased levels of PC, cardiolipin (CL), and coenzyme Q in mKO mitochondria, particularly in the soleus. Conversely, expression of CL biosynthetic enzymes was unchanged, and an *in vitro* binding assay indicated that STARD7 preferentially transfers linoleic acid-containing PC required for CL remodeling. Furthermore, electron microscopy revealed disorganized cristae structures, whereas 4-HNE-modified proteins, mtDNA content, and OPA1 processing remained unaffected. Together, these findings demonstrate that STARD7 plays an essential role for maintaining mitochondrial integrity and function in skeletal muscle, and its loss likely contributes to the pathogenesis of mitochondrial myopathy.

## Introduction

Cellular and organellar membranes are primarily composed of phospholipid bilayers. Among these, phosphatidylcholine (PC) is typically the most abundant phospholipid, accounting for more than 40% of total mitochondrial phospholipids and serving as a major structural component (Decker & Funai 2024). Mitochondria possess a double-membrane system comprising the outer mitochondrial membrane (OMM) and the inner mitochondrial membrane (IMM), both enriched in PC. Other mitochondrial phospholipids include phosphatidylethanolamine (PE, 25–35%), cardiolipin (CL, 5–15%), phosphatidylinositol (PI, 2–9%), phosphatidylserine (PS, ∼1%), and phosphatidylglycerol (PG, <1%). Mitochondria contain enzymes for the synthesis of PE, CL, and PG. In contrast, they lack the enzymatic machinery to synthesize PC, PI, and PS, which must therefore be supplied from other organelles such as the endoplasmic reticulum (ER) and Golgi apparatus.

Recent studies have shown that phospholipid transport into mitochondria does not occur through vesicular trafficking but rather through lipid transfer proteins (LTPs). These proteins shuttle lipids across membrane contact sites (MCSs) formed between mitochondria and other organelles. For instance, Mitofusin 2 not only establishes ER–mitochondria MCSs but also binds specifically to the phospholipid PS, thereby mediating its transfer from the ER to mitochondria (Hernandez-Alvarez *et al*. 2019). The OMM protein Mitoguardin-2 also forms ER–mitochondria MCSs and promotes the transfer of PS, PC, PE, phosphatidic acid (PA), and ceramides (Kim *et al*. 2022). Bridge-type LTPs such as VPS13 have recently been shown to mediate bulk phospholipid transport, including PC, across MCSs (Kumar *et al*. 2018; Adlakha *et al*. 2022). Despite these findings, the overall mechanisms of mitochondrial phospholipid trafficking remain incompletely understood.

STARD7 is an LTP that contains a mitochondrial targeting signal, a transmembrane domain, and a steroidogenic acute regulatory-related lipid transfer domain responsible for lipid transport. *In vitro* assays have demonstrated that STARD7 specifically transfers PC between liposomes (Horibata & Sugimoto 2010). The protein is localized to the outer leaflet of the OMM, to ER–mitochondria MCSs, and within the cytosol (Horibata *et al*. 2017). Loss of STARD7 leads to reduced mitochondrial PC levels and mitochondrial dysfunction, including abnormal cristae formation and decreased respiratory activity (Horibata *et al*. 2016; Yang *et al*. 2017; Uddin *et al*. 2024). Live-cell labeling analyses further revealed that STARD7 deficiency reduces PC delivery to mitochondria (Zhang *et al*. 2022). In addition, CRISPR screening using organelle-selective click labeling again identified STARD7 as a key factor in mitochondrial PC maintenance (Tsuchiya *et al*. 2023). Collectively, these studies indicate that STARD7 mediates PC transfer between the ER and mitochondria at MCSs, thereby supporting mitochondrial homeostasis.

Furthermore, STARD7 has been detected in the intermembrane space, where it transports PC between the OMM and IMM and thus contributes to CL biosynthesis (Saita *et al*. 2018). Recent findings also suggest that STARD7 has dual functions in coenzyme Q (CoQ) metabolism: mitochondrial STARD7 supports CoQ biosynthesis, whereas cytosolic STARD7 transfers CoQ to the plasma membrane to sustain antioxidant defense and ferroptosis resistance (Deshwal *et al*. 2023).

Mitochondrial dysfunction compromises energy metabolism and severely affects high energy–demanding tissues such as the brain, skeletal muscle, and heart. Disorders caused by mitochondrial DNA mutations, including MELAS, progressive external ophthalmoplegia, and Leigh syndrome, typically involve both muscular and neurological systems, resulting in myopathy and cardiomyopathy (Taylor & Turnbull 2005). Several rare inherited disorders are also linked to defects in mitochondrial phospholipid metabolism. Barth syndrome, for example, is a myopathy caused by mutations in *TAZ* (*TAFAZZIN*), which encodes a mitochondrial acyltransferase required for remodeling CL into its mature form. This disorder features reduced CL levels, impaired respiration, and abnormal cristae morphology (Schlame & Ren 2006; Schlame 2013). Other congenital diseases arise from mutations in upstream CL biosynthetic enzymes, including CDP-diacylglycerol synthase (CDS) –like protein TAMM41 and CL synthase 1 (CRLS1), causing cardiomyopathy and multisystem dysfunction (Lee *et al*. 2022; Thompson *et al*. 2022). The first step in PC biosynthesis is catalyzed by choline kinase, which forms phosphocholine. In skeletal and cardiac muscle, the β-isoform (CHKB) predominates. Loss of CHKB results in rostrocaudal muscular dystrophy (rmd), characterized by reduced mitochondrial PC, impaired respiratory complex activity, and hindlimb muscle weakness (Mitsuhashi *et al*. 2011; Mitsuhashi & Nishino 2011). Thus, defects in CL or PC biosynthesis cause myopathy or muscular dystrophy; however, the physiological effect of disrupted PC transport on skeletal muscle remains unclear. Previous reports showed that STARD7 deficiency in C2C12 myoblasts lowers mitochondrial PC, impairs respiration, and severely suppresses myotube formation, indicating a critical role for STARD7 in muscle physiology (Horibata *et al*. 2020; Rojas *et al*. 2024). However, the *in vivo* role in skeletal muscle and potential contribution to myopathy have not been established.

In this study, we examined the physiological function of STARD7 in skeletal muscle using muscle-specific knockout (mKO) mice. Loss of STARD7 disturbed mitochondrial phospholipid balance, leading to reduced respiratory function, altered cristae morphology, and markedly decreased endurance capacity. These results strongly suggest that STARD7 dysfunction contributes to the onset of mitochondrial myopathy.

## Materials and Methods

### Animals

All animal procedures were approved by the Committee for the Care and Use of Laboratory Animals at Dokkyo Medical University (Approval No. 1257). Embryos of C57BL/6NTac-*Stard7*^tm1a(EUCOMM)Wtsi^ were obtained from INFRAFRONTIER (Pettitt *et al*. 2009; Skarnes *et al*. 2011; Bradley *et al*. 2012; White *et al*. 2013). The C57BL/6J-FLPe mice (Kanki *et al*. 2006) were supplied by the RIKEN BRC through the National BioResource Project of MEXT/AMED, Japan. *Stard7*^tm1a^ mice were crossed with C57BL/6J-FLPe mice to generate *Stard7*^tm1c^ mice. Muscle-specific STARD7 knockout (STARD7-mKO) mice were produced by crossing *Stard7*^flox/flox^ mice with *Myf5*^Cre/+^ mice (strain No. 007893, The Jackson Laboratory, Bar Harbor, ME, USA) (Haldar *et al*. 2007). All mice were maintained on a C57BL/6N background for at least three generations and analyzed at ≤20 weeks of age.

### Immunoblot analysis

Skeletal muscle was lysed in 50 mM Tris-HCl buffer (pH 8.0) containing 8 M urea. Protein concentrations were determined using a BCA protein assay (Bio-Rad Laboratories, Hercules, CA, USA) after dilution with water. Equal amounts of protein (10 μg) were separated by SDS-PAGE and transferred to PVDF membranes (FluoroTrans; Pall Corp., Port Washington, NY, USA) using a Trans-Blot SD Semi-Dry Transfer blotter (Bio-Rad). Membranes were blocked with 5% (w/v) skim milk in TBS for 1 h, washed three times with TBS containing 0.1% Tween 20 (TBS-T), and incubated with primary antibodies overnight at 4°C. After three washes with TBS-T, membranes were incubated with horseradish peroxidase–conjugated secondary antibodies for 1 h at room temperature, washed again, and visualized with Clarity Western ECL Substrate (Bio-Rad) using a ChemiDoc Touch imaging system (Bio-Rad). Band intensities were quantified using Quantity One software (Bio-Rad), normalized to CBB staining, and analyzed within the linear detection range. Antibodies used were anti-STARD7 (15191-1-AP), UQCRC2 (14742-1-AP), PVALB (29312-1-AP), MYBPC2 (25257-1-AP), MCT4 (22787-1-AP), MYH4 (20140-1-AP), MYH7 (22280-1-AP), CRLS1 (14845-1-AP), and PGS1 (17149-1-AP) (Proteintech, Rosemont, IL, USA); anti-TAZ (sc-365810) (Santa Cruz Biotechnology, Dallas, TX, USA); OXPHOS complexes (ab110413) (Abcam, Cambridge, UK); anti-OPA1 (612606) (BD Biosciences, San Jose, CA, USA); and 4-HNE (MHN-100P) (JaICA, Shizuoka, Japan).

### Grip strength test

Muscle strength was measured using a grip strength meter (GPM-100B, Melquest, Japan). Mice were allowed to grasp a wire mesh with all four limbs, and their tails were gently pulled backward until release. The maximum tension recorded was considered one trial. Each mouse was tested twice, and the mean value was used for analysis.

### Exercise tolerance test

Endurance exercise capacity was evaluated using a motorized treadmill (MK680; Muromachi Kikai Co., Ltd., Tokyo, Japan), as previously described with slight modifications (Sato *et al*. 2015). After a 30 min acclimation period in the treadmill chamber, mice began running at 15 m/min on a 10% incline. An electrical stimulation plate at the rear of the treadmill encouraged continued running. The speed was increased by 1 m/min every minute for 10 min (up to 25 m/min). Exhaustion was defined as remaining at the rear of the treadmill for more than 5 s despite tail prodding.

### Measurement of blood lactate levels

Blood was collected from the tail vein immediately before and after 30 min of treadmill running at 25 m/min with a 10% incline. Blood lactate concentrations were determined using a Lactate Pro 2 meter (Arkray, Inc., Kyoto, Japan).

### Whole-transcriptome analysis by RNA-seq

Total RNA was extracted from the soleus muscle (three biological replicates per group) using a Maxwell RNC Simply RNA Tissue Kit (Promega Corp., Madison, WI, USA). mRNA libraries were prepared with a NEBNext Ultra II Directional RNA Library Prep Kit for Illumina, pooled at equal molar concentrations, and sequenced on an Illumina NextSeq500 using a 75-bp paired-end kit (Illumina Inc., San Diego, CA, USA). Sequencing reads were trimmed and mapped to the mouse GRCm38 release-94 reference genome using CLC Genomics Workbench (v22.0.1; Qiagen Inc., Germantown, MD, USA, RRID:SCR_011853). Read counts were normalized to transcripts per million (TPM), and log_2_ transformation was applied after adding 1 to all TPM values. RNA-seq data were analyzed using OlvTools (https://olvtools.com/).

### Preparation of muscle cryosections and immunofluorescence imaging

Muscle tissues were immediately embedded in cryomolds (Sakura Finetek, Torrance, CA, USA) filled with OCT compound (Sakura Finetek) and snap-frozen in isopentane chilled with liquid nitrogen. Sections (10 μm thick) were collected on glass slides (PRO-05; Matsunami Glass Ind., Ltd., Osaka, Japan) and stored at −30 °C until use. Acetone-fixed cryosections were rinsed in PBS containing 1% Tween 20 (PBS-T), washed, and blocked with blocking mouse IgG (MKB-2213-1; Vector Laboratories, Newark, CA, USA) and 5% normal goat serum (50062Z; Thermo Fisher Scientific, Waltham, MA, USA). The sections were then incubated overnight at 4°C with primary antibodies against myosin heavy chain (MHC) I (1:250), MHC IIa (1:250), MHC IIb (1:250) (BA-F8, SC-71, BF-F3; Developmental Studies Hybridoma Bank, University of Iowa, IA, USA), and laminin (1:500) (L9393; Sigma-Aldrich, St. Louis, MO, USA). The following day, sections were rinsed with PBS-T and incubated with secondary antibodies conjugated to Alexa Fluor 350, 488, 555, or 647 (A21140, A21042, A21127, A21244; Thermo Fisher Scientific). Slides were mounted with ProLong Gold Antifade Mountant (P36930; Thermo Fisher Scientific) and covered with glass coverslips. Fluorescent images were acquired using a DMi8 inverted microscope (Leica, Wetzlar, Germany).

### Isolation of mitochondria and measurement of oxygen consumption rate (OCR)

Mitochondria were isolated from skeletal muscle as described by Iuso *et al*. (Iuso *et al*. 2017). Muscle samples were minced in ice-cold mitochondrial isolation buffer (210 mM mannitol, 70 mM sucrose, 5 mM Hepes, 1 mM EGTA, 0.5% BSA, pH 7.2) and homogenized using a motor-driven Potter-Elvehjem homogenizer for 10 strokes at 600 rpm. The homogenate was centrifuged at 800 × g for 10 min, and the supernatant was further centrifuged at 8,000 × g for 10 min. The resulting mitochondrial pellet was resuspended in mitochondrial assay solution 1 (MAS1: 210 mM mannitol, 70 mM sucrose, 2 mM Hepes, 1 mM EGTA, 0.2% BSA, 10 mM KH_2_PO_4_, 5 mM MgCl_2_, pH 7.2). Protein concentration was determined by BCA assay (Thermo Fisher Scientific) after washing the pellet with MAS1 lacking BSA. OCR was measured using a Seahorse XFp Extracellular Flux Analyzer (Agilent Technologies, Santa Clara, CA, USA). Mitochondria (2 μg protein) were seeded onto a poly-L-lysine–coated plate (10 μg/ml) in MAS1 containing 10 mM pyruvate, 2 mM malate, and 4 μM FCCP. Sequential additions of 2 μM rotenone, 10 mM succinate, 4 μM antimycin A, and 10 mM ascorbate/0.1 mM TMPD were used to assess specific respiratory states.

### Blue Native PAGE

Blue Native PAGE was performed following established procedures (Wittig *et al*. 2006). Mitochondria were incubated with digitonin to solubilize membrane protein complexes under native conditions. After removal of insoluble material by centrifugation, Coomassie Brilliant Blue G-250 was added to the clarified extracts to stabilize the complexes. Samples were loaded onto 3–12% acrylamide gradient native gels and separated by electrophoresis. Following electrophoresis, proteins were transferred to PVDF membranes, and immunodetection was carried out using an anti-OXPHOS complexes antibody cocktail.

### Quantification of phospholipid molecular species and CoQ by LC–MS/MS

Mitochondria were isolated from skeletal muscle as described above, and lipids and CoQ were extracted with methanol containing internal standards: d_70_-PC (Olbracht Serdary Research Laboratories, Toronto, Canada); PA 14:0/14:0, PG 17:0/17:0, and CL 14:0 (Avanti Polar Lipids, Alabaster, AL, USA). Because CoQ was evaluated on a relative basis rather than as an absolute quantification, d_70_-PC was used as an internal standard for variability in extraction efficiency, injection volume, and MS response across samples. Phospholipids and CoQ were analyzed by reverse-phase HPLC using an L-column3 ODS (3 µm, 2.0 × 100 mm; CERI, Tokyo, Japan) coupled to a QTRAP 5500 mass spectrometer (Sciex, Framingham, MA, USA). The binary gradient consisted of solvent A (acetonitrile:methanol:water, 1:1:3, v/v/v, with 5 mM ammonium acetate) and solvent B (2-propanol with 5 mM ammonium acetate). The gradient program was 0–1 min, 95% A; 1–9 min, 5–95% B linear; and 9–13 min, 95% B. Flow rate was 0.2 ml/min, and column temperature was 40°C. Lipids and CoQ were quantified with MultiQuant v2.0 (Sciex), and normalized to internal standards and total protein.

### Specificity of STARD7 for PC molecular species

Recombinant 6×His-tagged STARD7 was expressed in *E. coli* as described previously (Horibata & Sugimoto 2010). Phospholipid vesicles containing 40 nmol of PC 16:0/18:1, PC 18:1/18:1, PC 16:0/18:2, or PC 18:0/20:5 (Avanti Polar Lipids) were prepared by sonication in 240 μl of 20 mM Tris-HCl buffer (pH 8.0) with 0.5 M NaCl. The reaction mixture containing STARD7 (1 nmol) and vesicles was incubated at 37°C for 10 min. The protein was recovered using TALON metal affinity resin (Takara Bio Inc., Shiga, Japan) and eluted with 20 mM Tris-HCl (pH 8.0) containing 250 mM imidazole. PC species bound to STARD7 were extracted with methanol and quantified by LC–MS/MS as described above.

### Electron microscopy

Soleus muscle samples were fixed in 2.5% glutaraldehyde, postfixed in 1% osmium tetroxide, and embedded in epoxy resin. Ultrathin sections were double-stained with uranyl acetate and lead citrate, and examined using a transmission electron microscope (HT7800; Hitachi High-Tech, Tokyo, Japan). For each mouse, ten fields were acquired at 5,000× magnification, and mitochondrial number and cross-sectional area were quantified in Fiji by manual tracing. Each field corresponded to an area of approximately 120 µm². Fields were randomly selected from non-overlapping areas of the soleus muscle to avoid sampling bias.

### Quantification of mitochondrial DNA (mtDNA)

mtDNA was quantified as described previously (Horibata *et al*. 2016). Total DNA was extracted using the standard proteinase K/phenol method. The copy numbers of mtDNA (*NADH dehydrogenase 1*) and nuclear DNA (*platelet endothelial cell adhesion molecule-1*, PECAM-1) were determined by quantitative PCR using PowerUp SYBR Green Master Mix (Thermo Fisher Scientific) on a QuantStudio3 instrument (Thermo Fisher Scientific).

## Statistical analyses

Quantitative data are presented as mean ± standard deviation (SD). Statistical analyses were conducted using GraphPad Prism 10 (GraphPad Software, San Diego, CA, USA). Comparisons between two groups were performed using unpaired two-tailed Student’s *t*-test or Welch’s *t*-test, and multiple-group comparisons by one-way ANOVA followed by Tukey’s post hoc test. Kaplan–Meier curves were compared using the log-rank test. A *p*-value < 0.05 was considered statistically significant, whereas *p* ≥ 0.05 was regarded as not significant (ns).

## Results

### STARD7 is highly expressed in mitochondria-rich slow-twitch muscle

Skeletal muscle fibers are broadly divided into fast-twitch and slow-twitch types. Fast-twitch fibers contract rapidly and support short bursts of activity, whereas slow-twitch fibers are rich in mitochondria, depend on oxidative metabolism, and play essential roles in sustained endurance exercise. In the mouse hindlimb, the gastrocnemius, tibialis anterior, and extensor digitorum longus are composed of fast-twitch fibers, while the soleus is enriched in slow-twitch fibers. Immunoblot analysis of these muscles showed that STARD7 was strongly expressed in the soleus, paralleling the mitochondrial marker UQCRC2 (ubiquinol-cytochrome c reductase core protein 2) (Fig. 1A, B). In contrast, the fast-twitch marker PVALB (parvalbumin) was detected in the gastrocnemius, tibialis anterior, and extensor digitorum. These findings suggest that STARD7 plays a major role in mitochondria-rich slow-twitch fibers, which are critical for maintaining endurance capacity.

**Figure 1.**
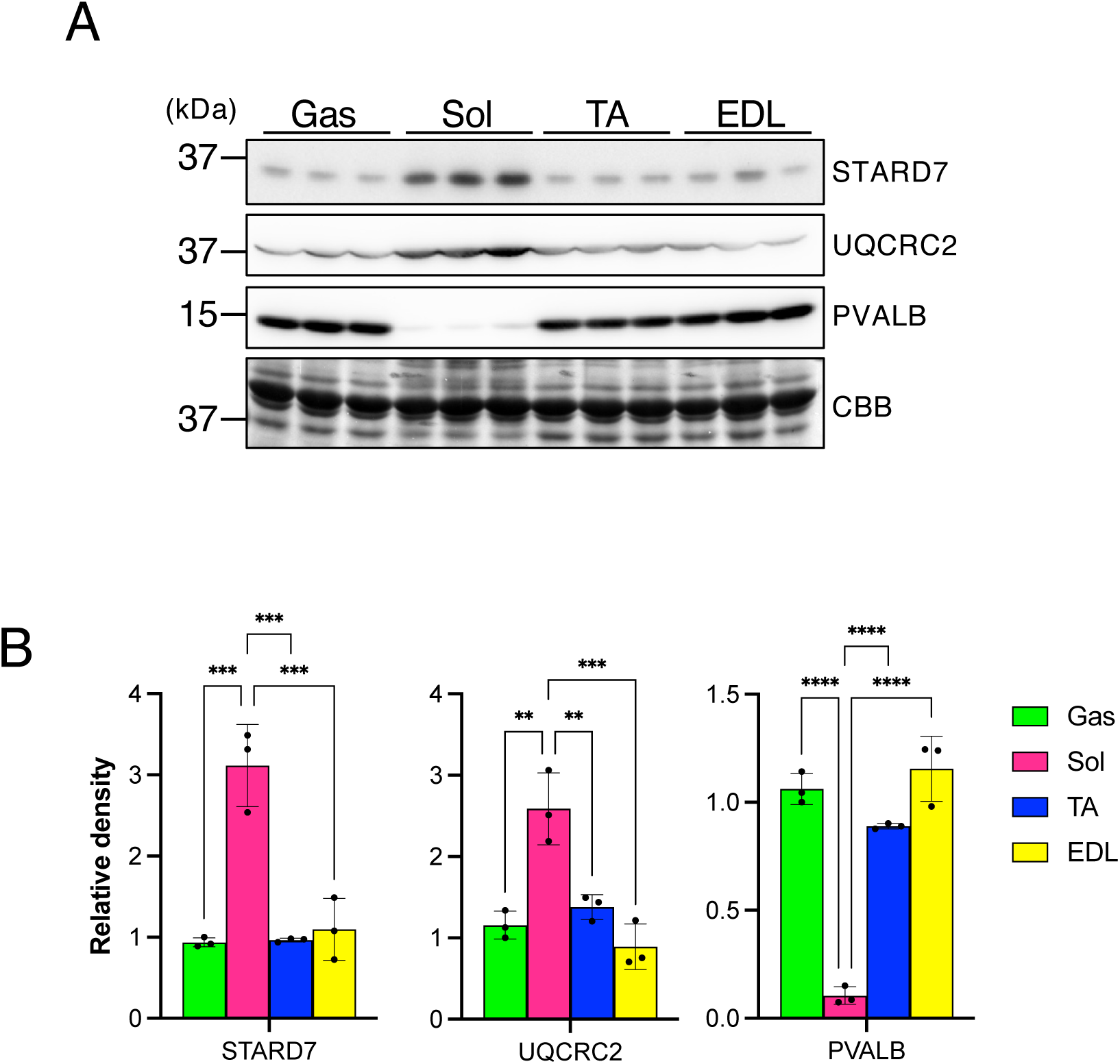
STARD7 is highly expressed in mitochondria-rich slow-twitch soleus muscle. (A) Expression of STARD7 in hindlimb muscles. UQCRC2 and PVALB serve as markers of mitochondria and fast-twitch fibers, respectively. CBB staining confirms equal protein loading. Each lane represents an independent animal (n = 3). Gas, gastrocnemius; Sol, soleus; TA, tibialis anterior; EDL, extensor digitorum longus. (B) Quantification of the immunoblot shown in (A). Statistical significance was determined by one-way ANOVA followed by Tukey’s post hoc test. **, ***, and **** indicate *p* < 0.01, *p* < 0.001, and *p* < 0.0001, respectively.

### Muscle-specific deletion of STARD7 impairs exercise performance without structural abnormalities

To clarify the role of STARD7 in skeletal muscle, *Stard7*^flox/flox^ mice were crossed with *Myf5-Cre* mice to generate skeletal muscle-specific knockout (mKO) mice. In these animals, recombination of *Stard7* exons 2 and 3 occurs in *Myf5*-expressing muscle progenitor cells around embryonic day 8 (Gensch *et al*. 2008). At 11 weeks of age, male mKO mice showed significantly lower body weight than controls, whereas female mice displayed no difference (Fig. 2A). Grip strength testing revealed reduced force in males but not females (Fig. 2B, C). Endurance capacity was next evaluated by treadmill running. Both male and female mKO mice exhibited markedly shortened running distances and durations, indicating decreased exercise endurance and metabolic capacity (Fig. 2D–F). Consistently, blood lactate levels measured before and after submaximal exercise were significantly elevated in the mKO group (combined male and female data) (Fig. 2G), indicating underlying metabolic dysfunction. To assess muscle structure, hematoxylin and eosin staining was performed on cross-sections, which revealed no pathological signs of myopathy, such as centrally nucleated fibers or intramuscular vacuoles. Together, these results indicate that loss of STARD7 impairs muscular performance, particularly endurance, without inducing structural abnormalities.

**Figure 2.**
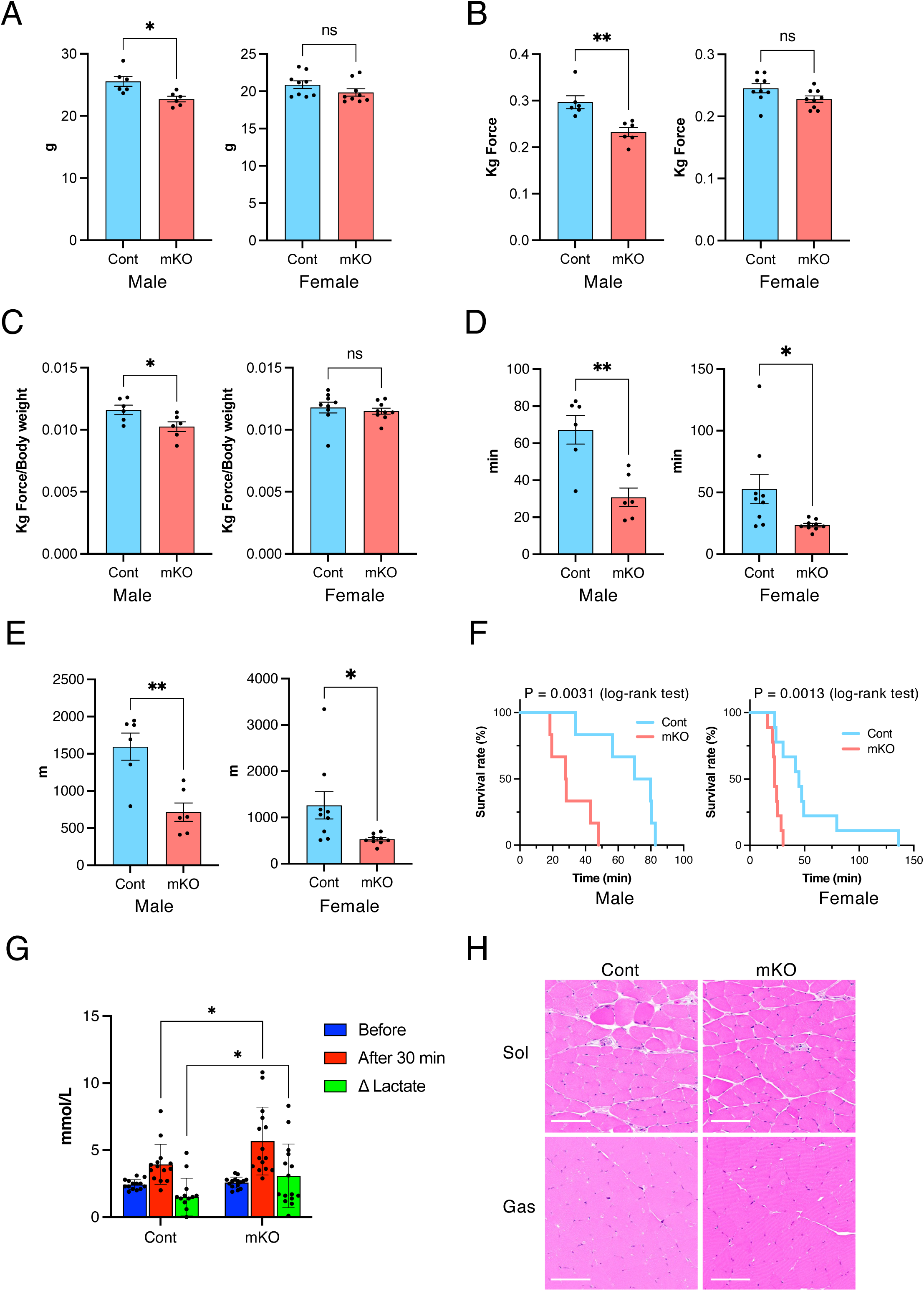
STARD7 deficiency reduces endurance capacity without structural muscle abnormalities (A) Body weight of 11-week-old mice. Cont = *Stard7* ^flox/flox^ control; mKO = muscle-specific knockout (*Myf5*-Cre; *Stard7*^flox/flox^). (B) Absolute forelimb grip strength (kilogram force). (C) Grip strength normalized to body weight (kilogram force/g). (D) Running time on a treadmill. (E) Running distance. For (A–E), male groups included six mice and female groups nine mice. Statistical significance was evaluated using unpaired two-tailed Student’s *t*-test. (F) Kaplan–Meier curve showing the proportion of mice remaining on the treadmill over time. Statistical significance was assessed using the log-rank test (male, n = 6; female, n = 9). (G) Blood lactate levels before and after fixed-intensity treadmill exercise. Data combine both sexes. Statistical significance was tested by unpaired two-tailed Welch’s *t*-test. * and ** indicate *p* < 0.05 and *p* < 0.01, respectively; ns, not significant. (H) Hematoxylin and eosin staining of cross-sections from the soleus (Sol) and gastrocnemius (Gas). Scale bar, 100 μm.

### Loss of STARD7 suppresses fast-twitch fiber–associated gene expression in the soleus

To explore the molecular basis of the reduced endurance in mKO mice, we conducted RNA-seq analysis of the soleus, where STARD7 expression is highest. Principal component analysis (PCA) revealed clear separation between control and mKO samples along principal component 1 (PC1), suggesting a broad transcriptional shift in the absence of STARD7 (Fig. 3A). A volcano plot of differentially expressed genes (Fig. 3B) showed significant downregulation of genes related to fast-twitch muscle structure and metabolism, including *Mybpc2*, *Actn3*, *Pvalb*, *Mct4*, and *Myh4*. In contrast, slow-twitch markers such as *Myh7* were unchanged. Heatmap clustering further distinguished control and mKO groups (Fig. 3C). Gene Ontology enrichment analysis identified “muscle system process” as the most enriched term (Fig. 3D). Consistent with these transcriptomic findings, immunoblot analysis confirmed reduced protein levels of MYBPC2, PVALB, MCT4, and MyHC-IIb (encoded by *Myh4*) in the soleus of mKO mice, whereas no significant changes were observed in the gastrocnemius (Fig. 3E). Expression of MyHC-I (encoded by *Myh7*) remained stable, indicating preserved slow-twitch fibers. Immunofluorescence analysis further showed that type IIb fibers, normally present in control soleus, were absent in mKO mice (Fig. 3F). These data demonstrate that in the soleus of STARD7-mKO mice, slow-twitch fibers are retained, while fast-twitch fibers are selectively reduced, altering overall fiber-type composition.

**Figure 3.**
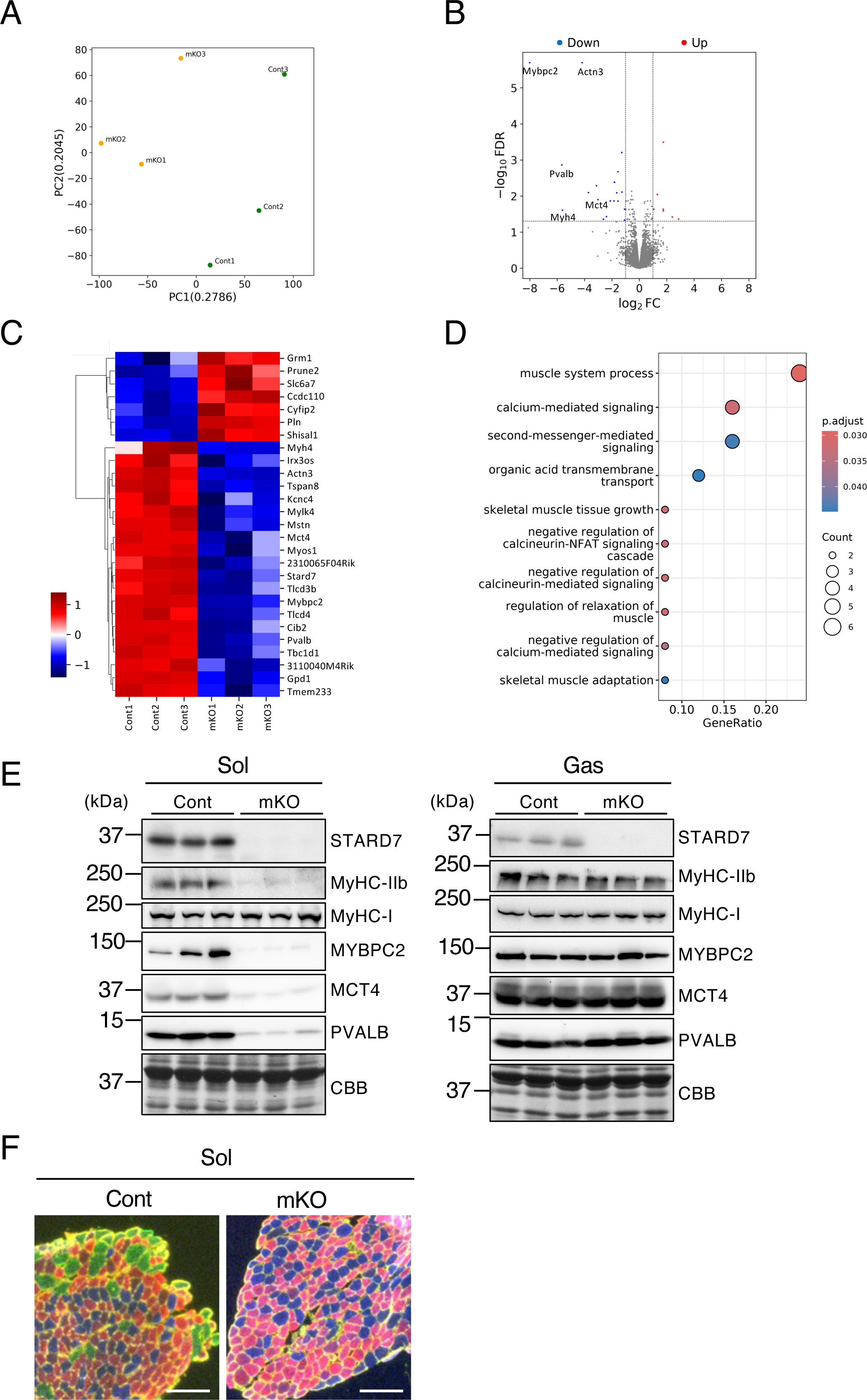
Loss of STARD7 disrupts fast-twitch–associated gene expression and fiber-type composition in soleus muscle (A) PCA of RNA-seq data from control (cont) and mKO soleus muscles. Each dot represents an individual animal (n = 3 per group). (B) Volcano plot of differentially expressed genes (DEGs). Differential expression was analyzed using the *edgeR* package after applying *filterByExpr* to remove low-expression genes. Genes with |log fold change| > 1 and FDR < 0.05 were defined as DEGs. Fast-twitch–related genes, including *Mybpc2*, *Actn3*, *Pvalb*, *Mct4*, and *Myh4*, were significantly downregulated in mKO. (C) Heatmap of DEGs (FDR < 0.05, |log fold change| > 1) in the soleus. (D) Functional enrichment analysis of DEGs using the *clusterProfiler* package for Gene Ontology (GO) and WikiPathways. GO terms and pathways with adjusted *p* < 0.05 were considered significantly enriched. (E) Immunoblot analysis of fast-twitch markers. Each lane represents an independent animal (n = 3). CBB staining confirms equal loading. (F) Representative image of soleus cross-sections stained with isoform-specific myosin heavy chain antibodies. Blue, type I (MyHC-I); red, type IIa (MyHC-IIa); green, type IIb (MyHC-IIb). Fibers lacking these isoforms were classified as type IIx. Scale bar, 200 μm.

### STARD7 deficiency reduces mitochondrial OCR

Although RNA-seq analysis revealed suppression of fast-twitch–associated genes, this alone could not explain the markedly reduced endurance capacity. Because mitochondrial function is crucial for endurance performance, we next measured mitochondrial OCR. Previous studies in cultured cells showed that STARD7 depletion diminishes mitochondrial oxygen consumption and disrupts mitochondrial function. To assess respiratory capacity *in vivo*, mitochondria were isolated from the soleus and gastrocnemius muscles, and OCR was measured using a Seahorse XFp flux analyzer after sequential addition of electron transport substrates and inhibitors. In both muscles, OCR driven by complex I substrates (pyruvate/malate) and by the complex II substrate (succinate) was consistently decreased (Fig. 4A, B). In contrast, OCR supported by complex IV substrates (ascorbate/TMPD) was reduced in the soleus but not in the gastrocnemius. To determine whether decreased OCR reflected instability of the respiratory complexes, representative subunits of complexes I–V were examined by immunoblot. Protein levels of these subunits were unchanged in both muscles (Fig. 4C), differing from earlier *in vitro* reports showing decreased MTCO1 (complex IV) upon STARD7 depletion (Horibata *et al*. 2016; Saita *et al*. 2018). Consistently, Blue Native PAGE analysis revealed no difference in respiratory supercomplex formation (Fig. 4D). These results indicate that the absence of STARD7 compromises mitochondrial electron transport activity without altering the expression or stability of respiratory complexes or their supercomplex organization.

**Figure 4.**
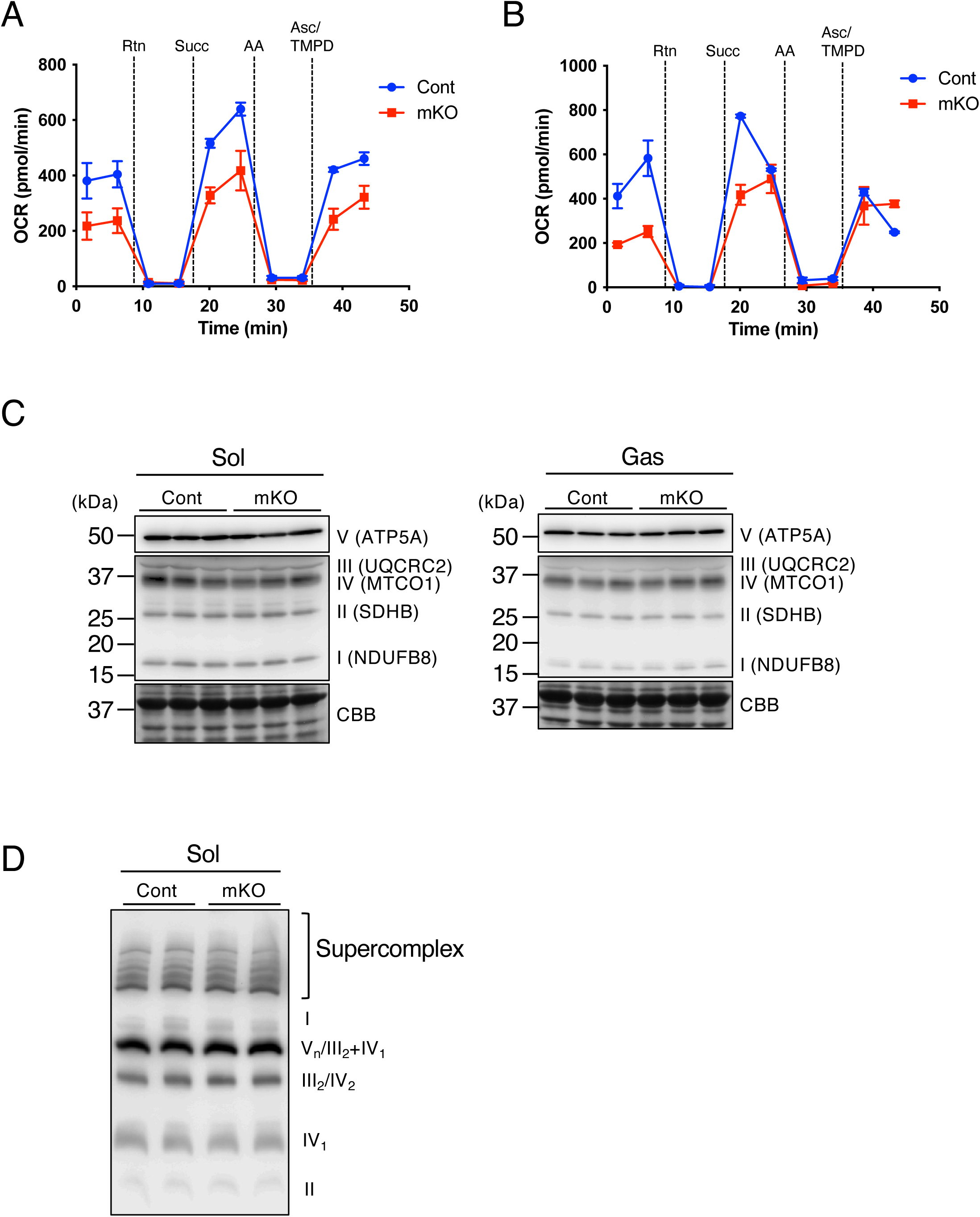
STARD7 deletion causes widespread defects in electron transport chain function independent of complex expression (A, B) Oxygen consumption rate (OCR) of mitochondria isolated from the soleus (A) and gastrocnemius (B) under various substrate and inhibitor conditions. Substrates included pyruvate/malate (complex I–driven), succinate (Succ; complex II–driven, in the presence of rotenone [Rtn] to inhibit complex I), and ascorbate/TMPD (Asc; complex IV–driven, in the presence of antimycin A [AA] to block upstream flow). Representative data from one of three independent experiments are shown. Similar results were obtained in all experiments. (C) Immunoblot analysis of representative subunits from mitochondrial respiratory complexes I–V. Data are from three independent animals. (D) Blue Native PAGE showing respiratory supercomplex formation. Shown are representative data, with similar outcomes in three independent experiments.

### STARD7 deficiency disrupts mitochondrial phospholipid homeostasis

Previous studies have established that STARD7 is crucial for maintaining mitochondrial PC. To determine whether this role is preserved *in vivo*, mitochondria were isolated from the soleus and gastrocnemius muscles, and their phospholipid composition was analyzed by LC–MS/MS. In the soleus, multiple PC molecular species were consistently reduced, indicating an overall decline in mitochondrial PC under STARD7 deficiency (Fig. 5A, B). In contrast, PC levels in gastrocnemius mitochondria were unchanged, suggesting a muscle type–specific dependency on STARD7 (Fig. 5C, D). Examination of CL revealed a marked reduction in both the soleus (Fig. 5E) and gastrocnemius (Fig. 5F). Because recent work showed that STARD7 participates in CoQ transport and biosynthesis (Deshwal *et al*. 2023), we further quantified mitochondrial CoQ levels. As shown in Fig. 5G (soleus) and Fig. 5H (gastrocnemius), the amounts of ubiquinone-9 (CoQ9) and ubiquinol-9 (CoQ9H_2_) were consistently reduced in both muscles. Collectively, these results demonstrate that STARD7 is required for maintaining CL and CoQ homeostasis in skeletal muscle mitochondria and plays a particularly important role in preserving PC content in the soleus.

**Figure 5.**
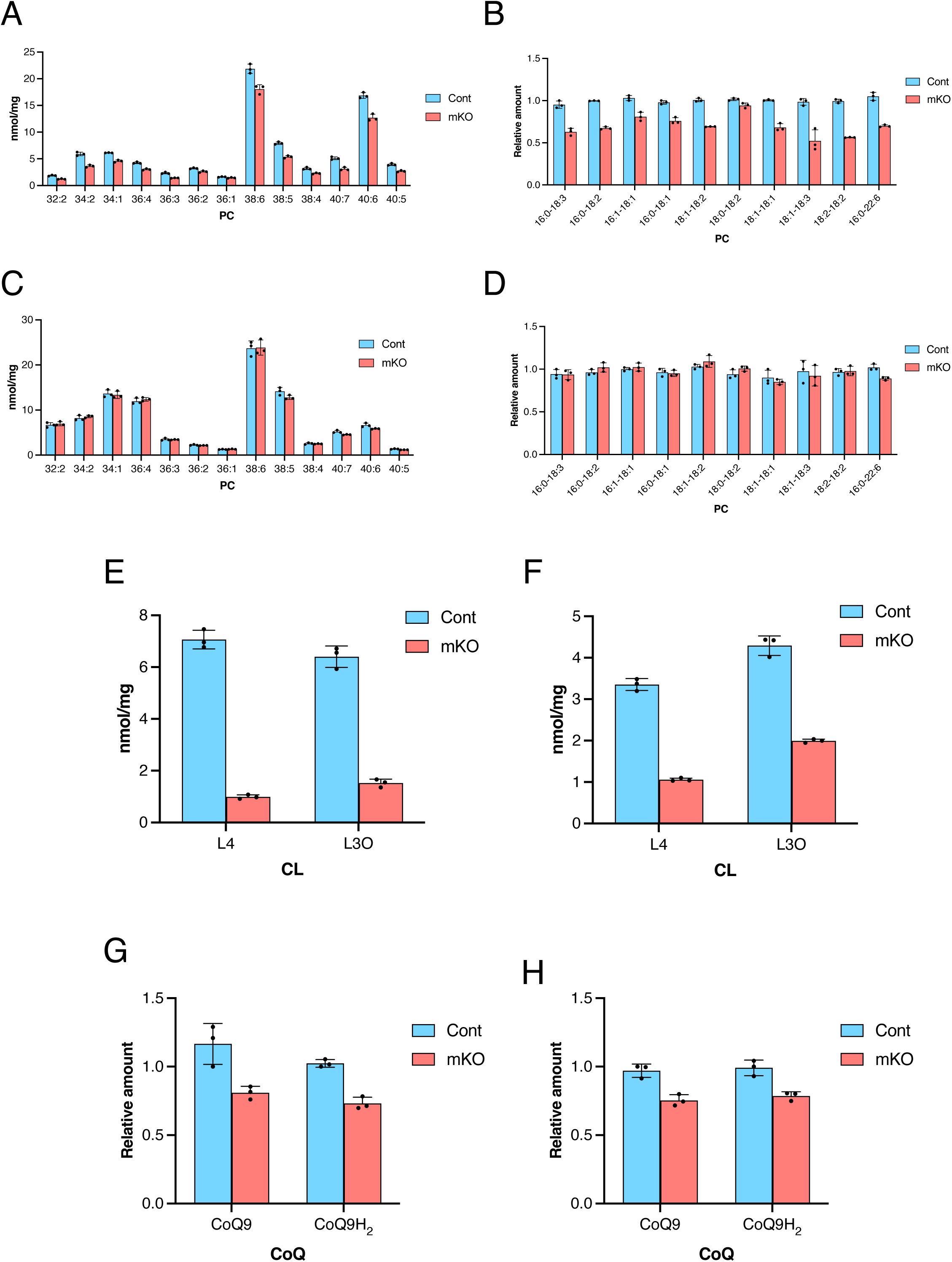
STARD7 deficiency disrupts mitochondrial PC, CL, and CoQ homeostasis in skeletal muscle (A) LC–MS/MS analysis of PC species in soleus mitochondria, acquired in positive ion mode and presented as total acyl chain composition (carbon number:double bonds). (B) Negative ion mode analysis of PC species in soleus mitochondria, resolving fatty acyl chains at the sn-1 and sn-2 positions. (C, D) PC species in gastrocnemius mitochondria analyzed under positive (C) and negative (D) ion modes. (E, F) CL levels in mitochondria from the soleus (E) and gastrocnemius (F). L4, tetralinoleoyl-CL; L3O, trilinoleoyl-oleoyl-CL. (G, H) CoQ levels in mitochondria from the soleus (G) and gastrocnemius (H). CoQ9, ubiquinone-9; CoQ9H_2_, ubiquinol-9. Representative data from one of three independent experiments are shown. Similar results were obtained in all experiments.

### Selective transport of linoleoyl-PC by STARD7 contributes to CL maintenance

CL is synthesized within the IMM through the pathway shown in Fig. 6A. To clarify the mechanism responsible for the reduced CL in mKO mice, we quantified phospholipid intermediates in the CL biosynthetic pathway. The levels of PA, a CL precursor, and phosphatidylglycerol (PG), an intermediate, were not reproducibly altered in either soleus or gastrocnemius (Fig. 6B–E). Immunoblot analysis further showed that protein levels of phosphatidylglycerophosphate synthase 1 (PGS1; the rate-limiting enzyme), cardiolipin synthase 1 (CRLS1; the enzyme catalyzing *de novo* synthesis), and TAFAZZIN (TAZ; the remodeling enzyme producing linoleoyl-enriched mature CL) remained unchanged in mKO muscles (Fig. 6F). These data indicate that reduced CL levels in mKO mice are not due to restricted substrate supply or loss of biosynthetic enzyme expression.

**Figure 6.**
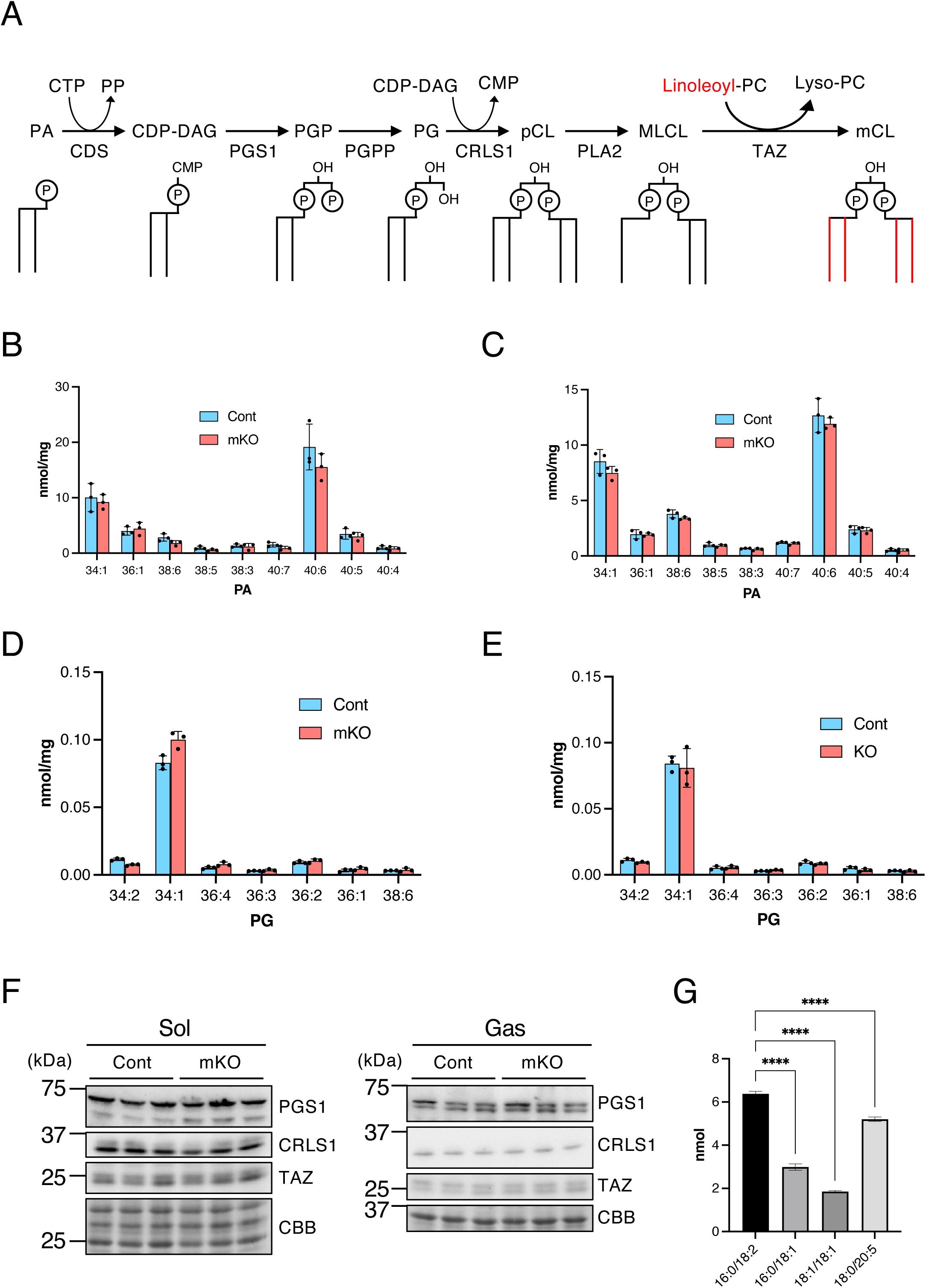
STARD7 supports CL homeostasis via selective transport of linoleoyl-PC (A) Schematic representation of the CL biosynthetic pathway. Linoleoyl-PC serves as a major fatty acid donor during TAFAZZIN (TAZ)-mediated CL remodeling. pCL, premature CL; MLCL, monolyso-CL; mCL, mature CL. (B–E) Quantification of PA (B, C) and PG (D, E) in mitochondria isolated from the soleus (B, D) and gastrocnemius (C, E) muscles. Shown are representative data, with similar results obtained in three independent experiments. (F) Expression levels of enzymes involved in CL biosynthesis. Data represent three independent animals. (G) Liposome binding assay assessing the fatty acid selectivity of STARD7. PC liposomes containing equal amounts of 16:0/18:2, 16:0/18:1, 18:1/18:1, and 18:0/20:5 species were incubated with recombinant STARD7, and PC species bound to the protein were quantified by LC–MS/MS. Statistical significance was evaluated using one-way ANOVA followed by Tukey’s post hoc test. **** indicates *p* < 0.001.

In skeletal muscle mitochondria, CL consists of linoleoyl-enriched species such as L4CL (tetralinoleoyl-CL) and L3OCL (trilinoleoyl-oleoyl-CL). Linoleic acid moieties in these forms are derived from linoleoyl-PC via TAZ-mediated remodeling. Previous work proposed that STARD7 contributes to CL biosynthesis by facilitating PC transfer from the OMM to the IMM (Saita *et al*. 2018). To evaluate whether STARD7 shows fatty acid selectivity in PC transport, recombinant STARD7 was incubated with liposomes containing equal amounts of four PC species (16:0/18:2, 16:0/18:1, 18:1/18:1, and 18:0/20:5), and bound PC species were analyzed by LC–MS/MS. As shown in Fig. 6G, STARD7 exhibited the highest affinity for 16:0/18:2 (linoleoyl-PC), followed by 18:0/20:5, 16:0/18:1, and 18:1/18:1. These findings suggest that, although a wide range of PC species is reduced in mKO mitochondria, STARD7 preferentially transfers linoleoyl-PC, thereby sustaining CL remodeling and maintaining CL levels in skeletal muscle mitochondria.

### Abnormal mitochondrial morphology in STARD7-deficient muscle

In cultured cells, the loss of STARD7 results in reduced and disorganized mitochondrial cristae. To assess these structural changes *in vivo*, we examined the soleus muscle of mKO mice by transmission electron microscopy. Mitochondria retained intact outer membranes but displayed highly disorganized inner membranes (Fig. 7A). No such abnormalities were observed in control animals. Quantitative analysis showed that mitochondrial number per area was unchanged (Fig. 7B), whereas mitochondrial cross-sectional area was significantly larger in mKO muscle (Fig. 7C). These findings indicate that STARD7 loss leads to disorganization of the inner membrane and mitochondrial enlargement without altering mitochondrial abundance.

**Figure 7.**
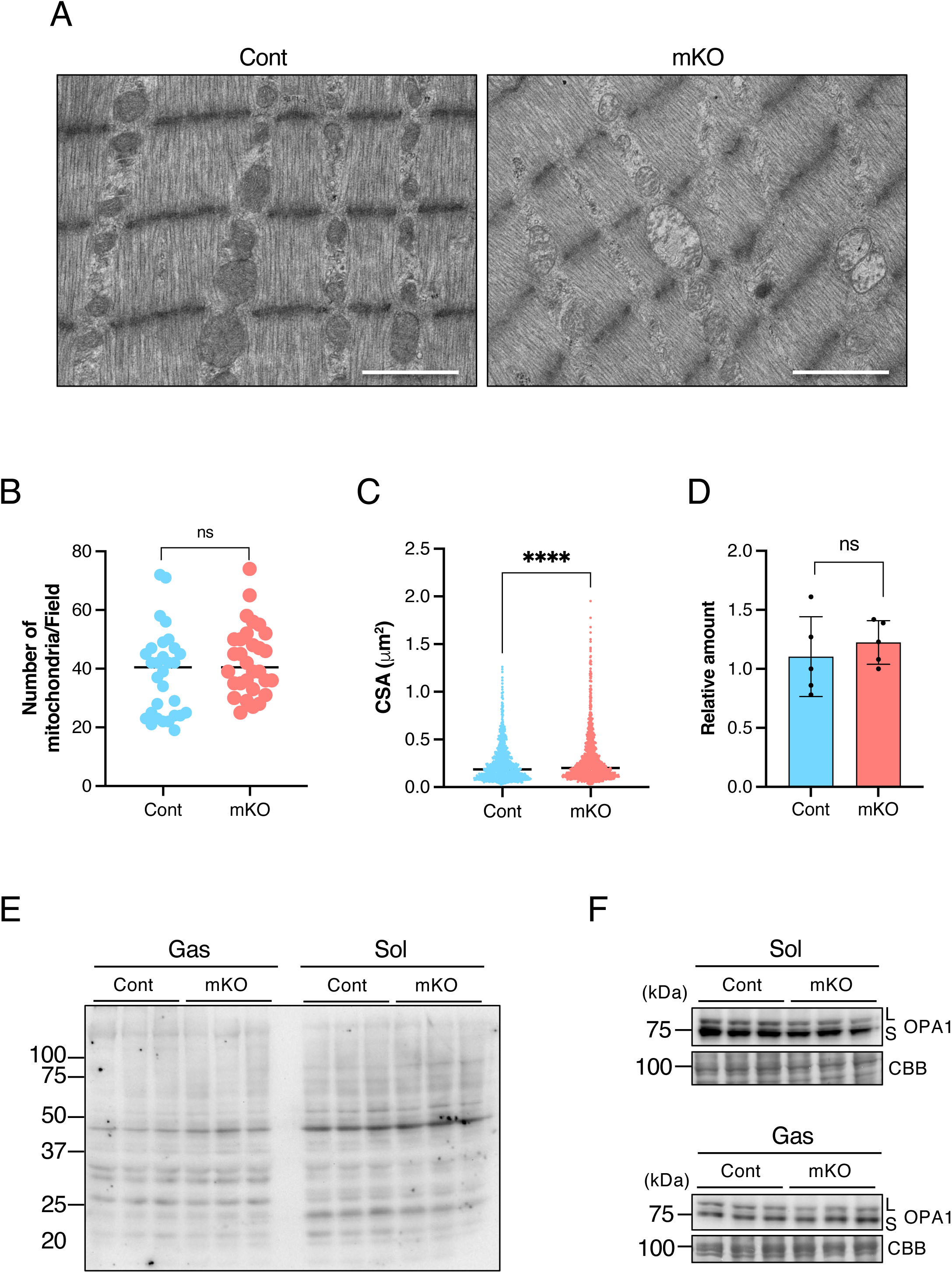
STARD7 deficiency causes abnormal cristae morphology without altering mtDNA content, increasing 4-HNE modification, or affecting OPA1 processing (A) Transmission electron micrographs of mitochondria in the soleus muscle. Scale bar, 1 μm. (B) Mitochondrial number per unit area in the soleus. Each dot represents one field. (C) Mitochondrial cross-sectional area in the soleus. Each dot represents one mitochondrion. (D) mtDNA copy number in the soleus muscle. Data were obtained from five mice per group. (E) Immunoblot analysis of 4-HNE–modified proteins. (F) Immunoblot analysis of OPA1 processing, showing membrane-bound long (L) and soluble short (S) forms. CBB staining confirms equal loading. Quantification (B, C) and immunoblotting (E, F) were performed using three independent animals (n = 3 per group). Statistical significance was assessed by two-tailed unpaired Welch’s *t*-test. **** indicates *p* < 0.001; ns, not significant.

In both Barth syndrome and rmd model mice, mitochondrial dysfunction is associated with elevated reactive oxygen species (ROS) production, increased 4-hydroxynonenal (4-HNE) modification, and reduced mitochondrial DNA (mtDNA) copy number (Mitsuhashi *et al*. 2011; Johnson *et al*. 2018; Suzuki-Hatano *et al*. 2019; Le *et al*. 2020). In contrast, no significant changes in mtDNA content (Fig. 7D) or 4-HNE modification (Fig. 7E) were detected in mKO mice. It has been reported that OPA1 processing is disrupted in Barth syndrome (Russo *et al*. 2024; Yazawa *et al*. 2025); however, in STARD7-mKO mice, the relative abundance of long (L) and short (S) OPA1 forms was comparable to controls, indicating normal OPA1 processing. Overall, these results suggest that although STARD7 deficiency impairs mitochondrial function and cristae organization, the pathological effects are milder than those observed in Barth syndrome or rmd models.

## Discussion

Although the role of STARD7 in mitochondrial phospholipid transport has been characterized in cultured cells, its physiological importance *in vivo* in skeletal muscle is not yet fully understood. This study sought to clarify how STARD7 maintains mitochondrial phospholipid balance and supports skeletal muscle function. In the soleus muscle of STARD7-mKO mice, we observed marked reductions in PC and CL, accompanied by abnormal mitochondrial structure, impaired respiration, and reduced endurance capacity. These findings demonstrate that STARD7 plays essential roles not only in preserving mitochondrial integrity but also in maintaining skeletal muscle performance.

STARD7 has been implicated in three key processes: (i) mediating PC transfer from the ER or Golgi to mitochondria (Horibata *et al*. 2017), (ii) sustaining CL biosynthesis by shuttling PC from the OMM to the IMM (Saita *et al*. 2018), and (iii) maintaining the CoQ pool required for efficient electron transport (Deshwal *et al*. 2023). In the present study, all three metabolites, PC, CL, and CoQ, were consistently reduced in the soleus muscle, where STARD7 expression is highest, indicating impairment of all three functions (Fig. 5). By contrast, the gastrocnemius muscle showed reductions in CL and CoQ but not in PC, suggesting that the second and third functions are predominantly affected in that tissue. CL maintains IMM curvature and provides the lipid microenvironment necessary for cytochrome c to mediate electron transfer between complexes III and IV. CoQ acts as a mobile electron carrier between complexes I/II and III. Therefore, simultaneous depletion of these metabolites likely destabilizes the IMM, resulting in impaired respiratory chain activity (Fig. 4), diminished ATP production, and reduced endurance capacity (Fig. 2).

Barth syndrome is the archetypal disorder linked to defective CL metabolism and arises from mutations in *TAZ*, which encodes an IMM-localized acyltransferase (Bione *et al*. 1996; Saric *et al*. 2015). TAZ catalyzes the transfer of linoleic acid from linoleoyl-PC to monolyso-CL (MLCL) and is indispensable for producing L4CL, the mature form of CL (Fig. 6A). In skeletal and cardiac muscle from Barth syndrome patients and model mice, reduced CL destabilizes respiratory supercomplexes, disrupts cristae morphology, and causes myopathy (McKenzie *et al*. 2006; Xu *et al*. 2016). Similarly, STARD7-mKO mice exhibited lower CL content and abnormal cristae. However, we detected no destabilization of respiratory complexes, elevated ROS, reduction in mtDNA, or altered OPA1 processing (Fig. 7), indicating that STARD7 deficiency causes a milder mitochondrial dysfunction. This milder phenotype likely reflects that STARD7 does not directly catalyze CL biosynthesis but instead provides linoleoyl-PC, the substrate for TAZ-mediated remodeling. Consequently, loss of STARD7 would limit CL remodeling without abolishing CL formation, resulting in moderate mitochondrial impairment. This mechanistic distinction may explain the milder phenotype of STARD7-mKO mice compared with Barth syndrome. To date, no skeletal muscle disorders have been reported in humans with *STARD7* mutations. Comparative analyses of STARD7 deficiency, Barth syndrome, and other models of impaired CL metabolism, including *CDS* and *CRLS1* deficiencies, will be critical to elucidate the connection between CL remodeling defects and myopathy pathogenesis.

A recent study identified a slow-twitch–specific super-enhancer that activates the *Stard7* promoter, driving its high expression in the porcine soleus (Tan *et al*. 2025). This may underlie the elevated *Stard7* expression observed in the mouse soleus (Fig. 1). The same study showed that transient siRNA-mediated STARD7 knockdown in porcine satellite cells or in mouse gastrocnemius muscle shifted fiber composition toward the fast type, reducing slow fibers. In contrast, constitutive deletion of STARD7 in the mouse soleus in the present study selectively decreased fast fibers while leaving slow fibers unaffected (Fig. 3). Thus, the consequences of STARD7 loss differ between transient and constitutive suppression. Notably, deletion of the mitochondrial fission factor DRP1 in C2C12 cells or in the mouse soleus also caused a reduction in fast fibers while preserving slow fibers (Yasuda *et al*. 2023). In that model, mitochondrial morphological abnormalities induced the stress cytokine GDF-15, which suppressed differentiation into fast fibers. Because mitochondrial structural defects were observed in STARD7-mKO muscle, a comparable stress-mediated mechanism may contribute to the selective loss of fast fibers. Future studies will be required to clarify how STARD7 deficiency influences mitochondrial stress signaling and fiber type regulation.

To date, three STARD7-deficient mouse models have been generated. In the global knockout, most embryos were nonviable, and the few surviving pups were not analyzed for skeletal muscle abnormalities (Yang *et al*. 2015). In contrast, epithelial-specific deletion in the lung caused mitochondrial dysfunction and barrier disruption (Yang *et al*. 2017), while intestinal epithelial knockout aggravated inflammation due to mitochondrial defects and impaired epithelial integrity (Uddin *et al*. 2024). Although PC and CL contents were not evaluated in these models, all exhibited mitochondrial structural and functional abnormalities leading to severe pathology. Together with our mKO results, these findings indicate that STARD7 is indispensable for mitochondrial integrity and function across multiple organs, and its loss produces distinct tissue-specific outcomes.

In summary, STARD7 is crucial for maintaining mitochondrial phospholipid homeostasis in skeletal muscle. Although overt dystrophic changes were not observed histologically, STARD7 deficiency reduced PC, CL, and CoQ levels, leading to mitochondrial dysfunction and diminished endurance capacity. These results provide new mechanistic insight into the development of a mild form of mitochondrial dysfunction associated with myopathy. Future studies identifying human *STARD7* variants and assessing their pathogenic relevance will be essential to establish the clinical significance of these findings.

## Data Availability

All numerical data supporting the figures will be provided as part of the Supplementary Information upon acceptance. RNA-seq data will be deposited in the NCBI Gene Expression Omnibus (GEO) and will be made publicly available upon acceptance.

## Author Contributions

Y.H.: Conceptualization, Methodology, Investigation, Formal Analysis, Data Curation, Visualization, Writing – Original Draft, Funding Acquisition

T.S.: Methodology, Investigation, Formal Analysis, Resources, Visualization M.O.: Methodology, Resources, Data Curation

S.Y.: Investigation, Visualization, Data Curation M.I.: Investigation, Supervision

S.M.: Resources, Supervision, Writing – Review & Editing A.K.: Supervision, Writing – Review & Editing

H.S.: Resources, Supervision, Writing – Review & Editing

## Acknowledgments

We thank Dr. Kinichi Matsuyama (Department of Pathology, Dokkyo Medical University Hospital) for valuable assistance with electron microscopy. We also acknowledge the Wellcome Trust Sanger Institute Mouse Genetics Project (Sanger MGP) and its funders for providing the mutant mouse line [EM:07272], and INFRAFRONTIER/EMMA (www.infrafrontier.eu) partner Biocenter Oulu, from which the mouse line was obtained. Information on project funding is available at www.sanger.ac.uk/mouseportal, and related phenotypic data are accessible at www.mousephenotype.org. This work was supported in part by JSPS KAKENHI Grant Numbers 24K10055 (Y.H.) and 24K02885 (S.M.), and by the Takeda Science Foundation (Y.H.).

